# Design and fabrication of flexible biodegradable microelectrode array for recording electrocorticography signals

**DOI:** 10.1101/2023.11.07.566132

**Authors:** Suman Chatterjee, Rathin K. Joshi, Sreenivas Bhaskara, RS Hari, Shubham Datta, KV Shabari Girishan, Hardik J. Pandya

## Abstract

Conventional flexible implantable microelectrode arrays, used for real-time brain activity monitoring, incur minimal tissue damage due to their flexibility; however, these require retrieval surgery, causing trauma to subjects. Implants, fabricated using biodegradable materials, dissolve gradually in body fluid, eliminating the necessity for retrieval surgery. This work reports designing and developing a 10-channel flexible biodegradable microelectrode array to acquire electrocorticography signals from rats’ brains. The array was fabricated using tungsten, a transient metal, and PLLA:PCL (80:20), a biodegradable composite polymer. The subdural implantation of the array on the somatosensory cortex of rats (n=3) facilitated electrical biopotentials acquisition using OpenBCI Cyton Boards. The baseline activities, the induced epileptic discharges after peripheral electrical stimulation, and the recovered baseline activities after antiepileptic drug administration were recorded from sedated rats. The chronic baselines and evoked activities were also acquired from awake rats. The gradual decline of chronic baselines was evident, suggesting a possible dissolution of electrodes. Furthermore, the time-frequency analyses demonstrate the differences between baselines and induced epileptic activities. The signals recorded from microelectrode arrays demonstrate their potential to monitor brain activities in a chronic study. Histology of the harvested vital organs from the euthanized rats confirmed minimal tissue damage due to implantation.

## 1. Introduction

Implantable devices are becoming popular for monitoring and diagnosing nervous system abnormalities, owing to the immense potential for the intracranial recording of neural activities. However, post-implantation studies show that the mechanical mismatch with brain tissues and the long-term exposure of tissues to abiotic materials may lead to inflammation. Though the flexible implants can help minimize the mechanical mismatch and thus reduce potential immunological responses, these implants need retrieval surgery after the intended period of study to avoid the formation of lesions of bacteria [1], which may lead to further trauma to the subject [2]. The fabricated devices using biodegradable materials degrade after implantation (*in vivo* condition) depending on the ambient pH, temperature, and concentration of ions in the medium [3]–[5]. These devices can be designed to utilize for a predetermined period by choosing suitable materials and tuning their geometry. Thus, the flexible biodegradable implants show minimized inflammation due to their flexibility and excellent biocompatibility. They provide an escape to the retrieval surgery owing to the complete biodegradability of the constituent materials, compared to conventional permanent implants. Biodegradable materials include metals (e.g., magnesium, tungsten, molybdenum, etc.) [1], [4], insulators (e.g., silicon dioxide, silicon nitride, etc.) [2], [4], polymers (e.g., Polycaprolactone (PCL), poly(lactic acid) (PLA), poly(glycolic acid) (PGA), silk, etc.) and polymer composites (e.g., Poly(lactic-co-glycolic acid) (PLGA), poly(L-lactic acid): polycaprolactone (PLLA:PCL), etc.) [1], [3], [5] of diverse physical and chemical properties which offer development of complete biodegradable for a wide range of applications.

Epilepsy is one of the major neurological disorders which can be diagnosed by continuous monitoring of neural electrical activity. Moreover, localization of the epileptic lesion requires a high-resolution mapping of the cortical surface by an implantable high-density electrode array. Researchers have developed flexible biodegradable electrode arrays to record the electrical response of the brain for studying different animal models of epilepsy [1], [5]. The competence of the fabricated arrays to record Electrocorticography (ECoG) signals was demonstrated using *in vivo* experiments on animal models. ECoG signals can be recorded from the brain surface (for example, the motor cortex [6] and the somatosensory cortex [7], [8]) following implantation in animal models, such as mice [9], [10], rats [5], [11], [12], monkeys [13], [14]. The epileptic activities can be induced by the application of convulsant, for example, bicuculline [5], penicillin [1], [11], pentylenetetrazole [15], etc., or by electrical stimulation [7] in animal models.

The external electrical stimulation protocols do less harm to subjects and ensure better control over induced epilepsy than the chemically-induced methods [16], [17]. However, a common limitation of electrically induced seizures is seizure generalization in a large area rather than focal seizures, where seizures are localized in a specific brain region. On the contrary, the electrical stimulation applied directly to the brain causes damage to the tissue [7]. These evident the application of peripheral nervous stimulation as one of the preferred methods of eliciting epilepsy.

Biodegradable materials are utilized in fabricating partially and completely biodegradable arrays. In 2010, Kim et al. reported neural mapping of the brain surface by a partially biodegradable array using silk, a substrate-hardening material that gets dissolved and facilitates conformal contact for the electrode array [18]. This study recorded the visually evoked P100 response and sleep spindles from a feline brain. In 2014, Kozai et al. recorded chronic tissue responses using a carboxymethylcellulose-coated probe. The biodegradable carboxymethylcellulose was used to provide mechanical stiffness to the non-biodegradable probe [19]. Khilwani et al. reported a platinum neural probe encapsulated with bioresorbable Parylene-C dissolves, leaving behind the implanted electrodes [20]. In another report on a polyimide-based electrode array for neural recording, acid-terminated PLGA was used as an insertion device [21]. In these reports, the electrode array was not biodegradable; however, there are reports on completely biodegradable arrays for recording brain signals. In chronic studies, Yu et al. used a bioresorbable electrode array to record pre-ictal, ictal, and inter-ictal activities and sleep spindles from the cortical surface of rats. PLGA, a resorbable polymer, and silicon dioxide were used as substrate and encapsulation layer, respectively, and a doped silicon nanomembrane was used as recording electrodes. The four-channel fabricated array was immersed in PBS (pH 10) to study the dissolution, and it was dissolved in cerebrospinal fluid in ∼33 days in the chronic *in vivo* experiment [5]. In 2019, Xu et al. reported a biodegradable electrode array integrated with a pressure sensor to study dynamic changes in brain signals and intraoperative swelling of the cortex. The array was fabricated on a PLLA:PCL (80:20) substrate using molybdenum electrodes to record electrocorticography signals and a pressure sensor to measure intracranial pressure. The implanted device could effectively work for five days [1].

PLLA and PCL are more hydrophobic than other polymers or composites (for example, PGA, PLGA, etc.). The dissolution properties of the composite of PLLA and PCL are tuneable depending on the ratio of the polymers (%w/w). Thus, the PLLA:PCL is chosen to elongate the lifetime of the implant. Moreover, the flexibility of the PCL will contribute to increasing the overall flexibility of the composite film to achieve the desired mechanical properties [3].

To our knowledge, there are only a few studies on completely biodegradable microelectrode arrays for recording ECoG signals under different neurophysiological conditions. However, the effect of antiepileptic drugs (AEDs) during seizures on brain signals has not been studied using implantable biodegradable flexible arrays. Moreover, the topical application of a convulsant may cause a direct insult to the brain, and the chemically induced epilepsy may affect other organs along with the brain. Hence, peripheral electrical stimulation can elicit epilepsy to avoid organ damage and minimize brain tissue damage. This work reports the design and development of a novel, biodegradable, flexible microelectrode array (MEA) for a simultaneous 10-channel recording of ECoG signals from the somatosensory cortex of three rats. Due to its flexibility, the fabricated biodegradable MEA makes conformal contact with the cortical surface. The epileptic activities were induced by peripheral electrical stimulation after surgery under ketamine anesthesia, which was not reported earlier. Two OpenBCI Cyton Boards were used for ECoG signal acquisition. A schematic of the *in vivo* study is shown in Figure 1. The chronic *in vivo* experiments were performed to demonstrate that the fabricated MEA can record: (i) baselines, (ii) induced epileptic activities, (iii) recovered ECoG signals, and (iv) evoked responses (by touching whiskers) from the somatosensory cortex of a rats. The minimal tissue damage obtained by histology studies and induction of epileptic activities demonstrates the application of peripheral nervous stimulation as one of the preferred methods of eliciting epilepsy. This work also demonstrates the capability of the OpenBCI Cyton Boards for signal acquisition to provide a portable affordable alternate solution for *in vivo* biopotential acquisition.

**Figure 1.**
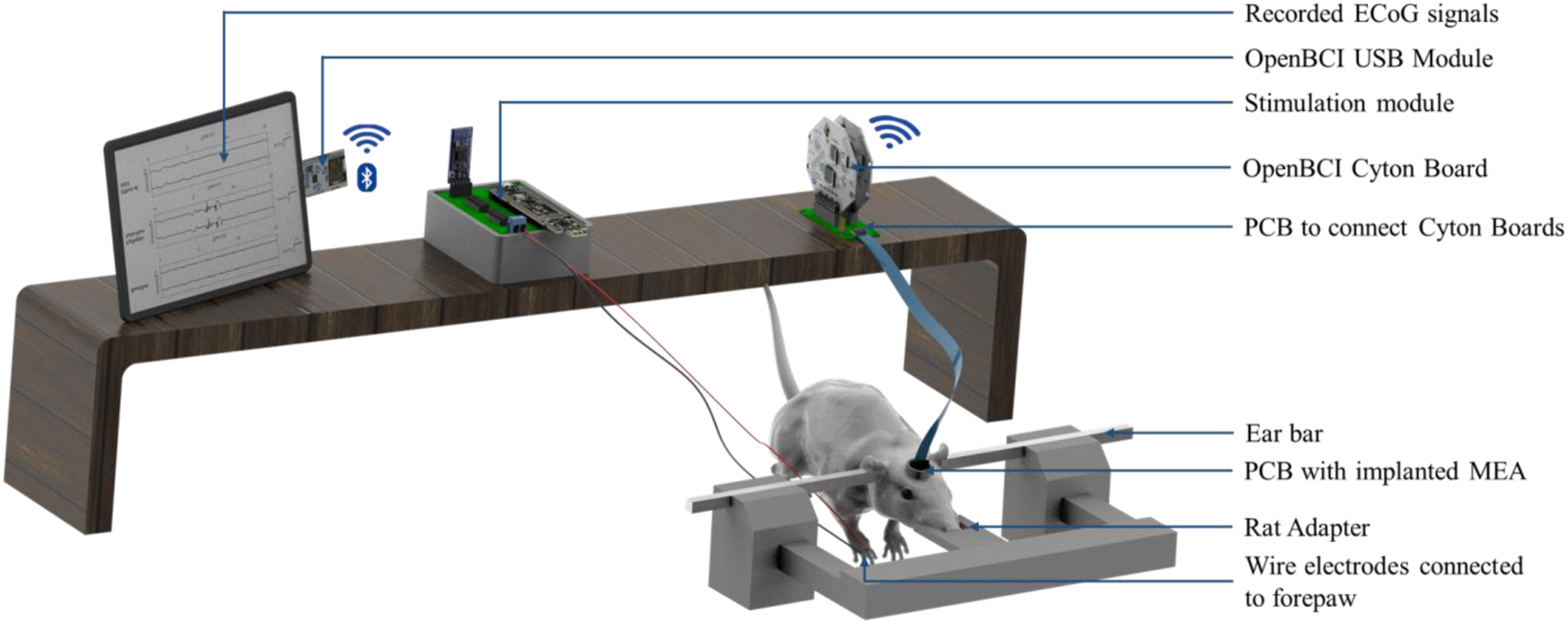
Illustration of the experimental setup for stimulating forepaw with wire electrodes (with a needle) and recording electrocorticography signals from sedated rat using a flexible, biodegradable microelectrode array.

## 2. Materials and methods

### 2.1 Design and fabrication of biodegradable microelectrode array

The biodegradable MEA with ten electrodes was designed with a diameter of 300 µm and a pitch of 1000 µm (center to center) (Figure 2(a)). The recording area of 3.3 mm × 2.04 mm was placed on the somatosensory cortex to study electrically-evoked seizures by peripheral stimulation.

**Figure 2.**
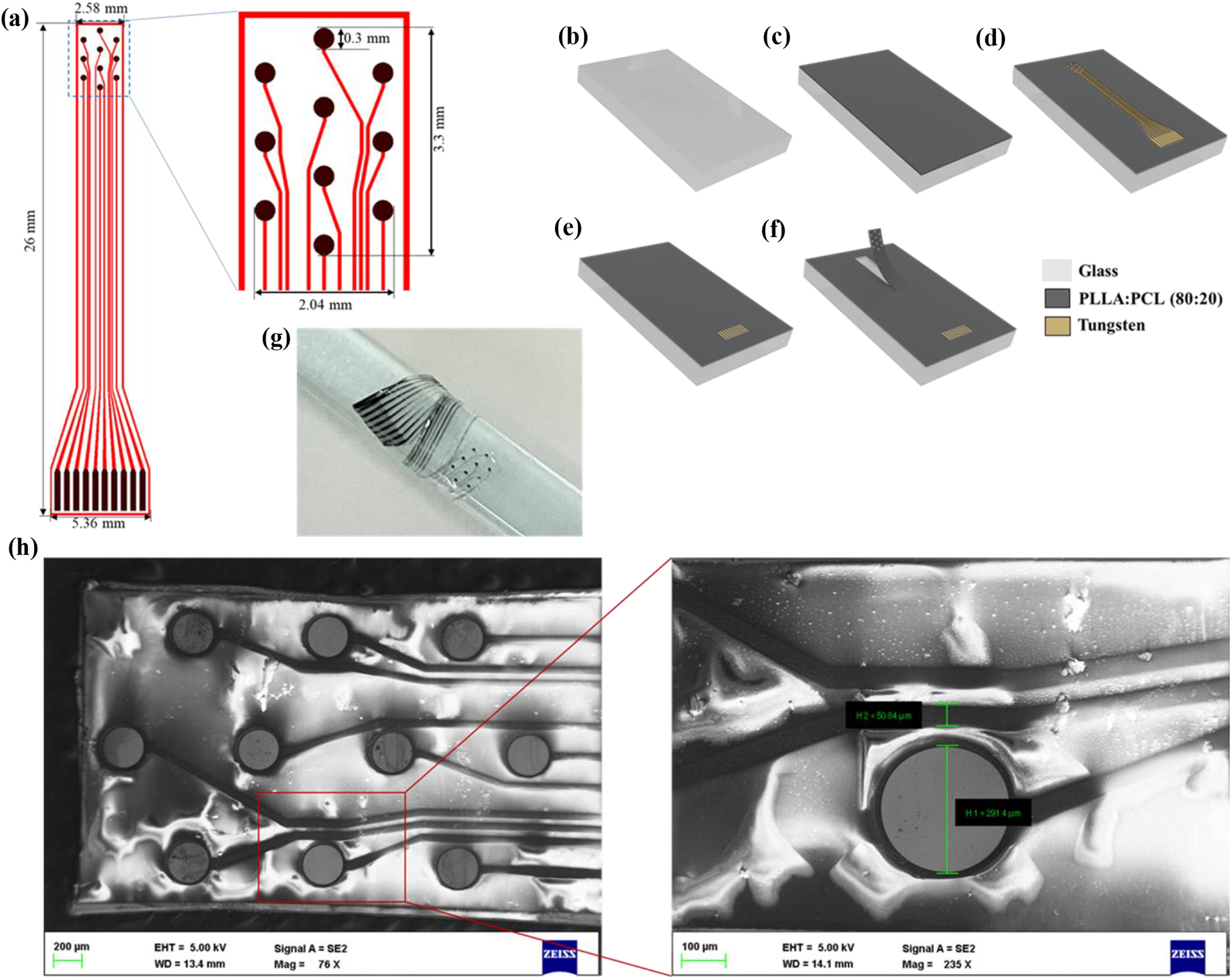
Biodegradable microelectrode array (MEA) for recording ECoG signals. **(a)** Schematic of the MEA (inset) the recording electrode array; **(b) – (f)** Process flow for fabricating the flexible biodegradable MEA: **(b)** Glass wafer, **(c)** Spin-coating of PLLA:PCL (80:20) solution in dioxane to form a film, **(d)** Metal (Tungsten of thickness 250 nm) deposition through shadow (metal) mask, **(e)** Spin-coating of PLLA:PCL (80:20) solution to form an insulation layer, followed by patterning for opening windows at the recording electrodes and the contact pads, **(f)** Plasma etching of polymer film to define the boundary followed by peeling off the fabricated microelectrode array from the glass substrate; **(g)** Photograph of the fabricated MEA, wrapped around a glass cylinder (diameter 6.4 mm), and **(h)** SEM image of the recording electrode array.

The MEA was fabricated using the reported biodegradable materials to achieve biodegradability after implantation. PLLA:PCL (80:20), a biodegradable polymer, was used as the substrate and the encapsulating material. The recording electrodes, contact pads, and their interconnecting metal lines were fabricated by tungsten, a transient metal. The composite polymer was prepared using poly(L-lactide) (average Mn 40000) (from Aldrich Chemistry) and polycaprolactone (average Mn 80000) (from Sigma-Aldrich), with a ratio of 4:1 (w/w), respectively. These polymers were dissolved in 1,4-Dioxane (anhydrous, 99.8%) (from Sigma-Aldrich) by continuous stirring at room temperature using a magnetic stirrer.

A four-inch glass wafer was cleaned by the Piranha cleaning process. A composite polymer PLLA:PCL (80:20) was spin-coated at 500 rpm for 1 minute on the clean glass wafer by dissolving in dioxane solution and was desiccated for 72 hours to prepare a film. This film acted as the substrate, and the glass wafer was used to support the polymer films during fabrication. A tungsten film of 250 nm thickness was deposited through a metal (shadow) mask by DC sputtering technique (*Tecport Optics, USA*) to realize the recording electrodes, contact pads, and interconnects. Then an encapsulation layer of PLLA:PCL (80:20) was deposited for electrical insulation and to protect the metal layer *in vivo*. This encapsulation was deposited by spin-coating at 1200 rpm for 1 minute, followed by desiccation for 72 hours. The encapsulation layer was patterned in a plasma chamber to open windows on the electrodes and contact pads. The PLLA:PCL layers were etched in a plasma chamber to define the MEA boundary, followed by peeling off from the glass wafer. The process flow of the MEA fabrication is illustrated in Figure 2(b) – (f).

The photograph of the fabricated MEA (wrapped around a glass cylinder with a diameter of 6.4 mm) and the SEM image of its recording area are shown in Figure 2(g) and Figure 2(h), respectively.

### 2.2 Characterization of the fabricated MEA

#### 2.2.1 Impedance Spectroscopy

The fabricated biodegradable microelectrode array was characterized by impedance spectroscopy for assessing the performance of the electrodes before implantation. Impedance spectroscopy was performed over a frequency range of 20 Hz to 10 kHz by immersing the recording electrode array in a phosphate buffer solution (PBS) of pH 7.4. The impedance was measured at 100 mV trigger over this frequency range as 200 data points using a commercial LCR meter (*LCR – 8105G*, *GWINSTEK*). The impedances of the consecutive electrode pairs were measured and plotted, and the average impedance of an electrode was determined at 1 kHz frequency.

#### 2.2.2 Bending test

Due to flexibility, cracks may occur in the patterned metal layer or the substrate and encapsulation layers of the fabricated MEA. The reliability of the array during a chronic *in vivo* study was evaluated by a bending test. The MEA was attached to a custom-designed PCB and mounted onto a micromanipulator indentation stage. The recording electrodes were immersed in PBS (pH 7.4). The impedance of five pairs of equidistant electrodes from the array was measured at a 1 kHz frequency with a 100 mV trigger when the device was bent by 60° because of the periodic vertical movements of the micromanipulator. The impedance was measured up to 500 cycles at an interval of 100 cycles to study the change in impedance of the electrodes due to bending.

### 2.3 Dissolution of the MEA

The dissolution study of the fabricated MEA was performed in phosphate buffer solution (PBS) prior to the implantation in a rat. The biodegradability of the fabricated array was tested by its degradation study after immersing it entirely in PBS (pH 7.4). The PBS with immersed MEA was stored at 37 °C and was continuously agitated using a shaker platform. The dissolution of the MEA was monitored at an interval of one hour for twenty-two hours. This dissolution study showed the degradation of the constituent materials in a bio-mimicked environment at the standard physiological temperature.

### 2.4 Impedance measurement of MEA during *in vitro* degradation

The *in vivo* degradation of an implanted MEA can be extrapolated from its *in vitro* degradation in PBS (pH 7.4). The change in impedance was observed by performing electrical impedance spectroscopy after immersing the recording electrode array in PBS. This can provide an idea of post-implantation impedance change due to encapsulation and patterned metal layer degradation. A fabricated MEA was attached to a PCB, and the impedance of consecutive electrode pairs was measured at a 1 kHz frequency with a 100 mV trigger. The impedance was measured at one hour interval until the recording electrodes were dissolved. The average impedance of the consecutive electrode pairs was plotted to understand the changes in impedance due to degradation.

As we do not have access to imaging techniques for rodents (such as animal MRI) for visualizing the *in vivo* degradation of the implanted MEA, the change in impedance of the electrodes during *in vitro* degradation was studied and was extrapolated to understand the *in vivo* degradation by the change in impedance resulting in decaying of recorded ECoG baselines.

### 2.5 Design and fabrication of interfacing PCBs

The PCBs were designed and fabricated to interface the MEA with the OpenBCI Cyton Boards for ECoG acquisition. The designed PCB used to attach the MEA has a dimension of 13 mm × 14 mm × 0.8 mm (l × w × h) and a weight of 0.32 g. The MEA was attached using a 10-pin FFC/FPC connector. This PCB has 10 data channels and two ground and reference pins for interfacing OpenBCI Cyton Boards. During the chronic study, this PCB was fixed on the rat’s skull using dental cement. Another PCB of dimension 60 mm × 28 mm × 0.8 mm (l × w × h) was designed and fabricated as an adapter to interface the acquisition system to the input data channels. The second PCB helps to minimize the weight on the rat’s head.

### 2.6 Design and implementation of stimulation module

The epileptic activities in rats were induced by peripheral electrical stimulation. A biphasic stimulus was applied to the forepaw of a rat to induce reflex seizures without damaging the brain [7]. A module stimulating the forepaw was designed and realized using a PSoC^®^ 5LP (*Cypress Semiconductor Corporation, USA*) microcontroller. A custom code was written in Embedded C to implement the stimulation cycle. The stimulation cycle of 70 s was divided into two segments. The first segment (35 s) includes the stimulation, and the second segment (35 s) is a relaxation period. The first segment was further divided into three sections, i.e., (i) 5 s of ECoG recording, (ii) 10 s of biphasic stimulation, and (iii) 20 s of ECoG recording to study the evoked responses. The 10 s stimulation section stimulated the forepaw with a rectangular wave (On-time: 1 ms and Off-time: 124 ms) of 8 Hz frequency and 2 mA amplitude. This stimulation cycle of 70 s was run ten times in each trial for ten trials per experiment to study the evoked seizures. This stimulation protocol elicited seizure activities in rats without direct chemical or mechanical insult to the brain.

The stimulation was applied by clinically validated pre-sterilized Medtronic DSN2280 Dual Electrodes *(Medtronic PLC, USA*) with 27G subdermal needle electrodes (stainless steel, 0.42 mm diameter), connected using extension lead wires. These needle electrodes deliver signals from the implemented stimulation module to the forepaw for eliciting epileptiform discharges by stimulating the peripheral nervous system.

### 2.7 *in vivo* studies

The chronic *in vivo* studies were performed on healthy male Sprague-Dawley rats (n = 3) of age 13 – 15 weeks and weight 350 g – 380 g. These chronic *in vivo* studies were performed following the Institutional Animal Ethics Committee guidelines (IAEC Project Number: CAF/Ethics/667/2019) of the Indian Institute of Science (IISc), Bangalore and Central Animal Facility (CAF) at IISc is registered with the Committee for the Purpose of Control and Supervision of Experiments on Animals (CPCSEA), Ministry of Environment and Forests, Government of India, New Delhi (Registration Number: 48/GO/Re-SL/Bi-S/99/CPCSEA, 11-03-1999). All methods are reported in accordance with ARRIVE guidelines (https://arriveguidelines.org). The rat was anesthetized by intraperitoneal administration of ketamine and xylazine (Dose: 80 mg/kg body weight and 10 mg/kg body weight, respectively). The anesthetized rat was fixed in a stereotaxic frame to arrest the movement during surgery. The surgical area was cleaned using Povidone-iodine. A curvilinear midline incision was performed following the positioning of the rat in the frame, and the skin flap was raised along with the periosteum. The exposed skull was cleaned with saline-soaked Q-tips, and the bleeding was stopped by electrocautery. The Bregma and Lambda points on the skull were identified, and a 5 mm × 4 mm craniotomy was performed to expose the somatosensory cortex forelimb region (S1FL) of the right hemisphere (Figure 3(a)). Two stainless steel screws were drilled into the skull above the left hemisphere and were used as the ground and the reference electrodes. The bone flap was elevated from the underlying dura and dipped in saline for placing back while sealing the craniotomy defect. The dura incision was performed posteriorly and raised as a flap. The fabricated array was implanted subdurally. The bone flap was placed back on the craniotomy defect and was sealed using dental cement. After the surgery, the rat was allowed to come out of anesthesia and recover in a cage.

**Figure 3.**
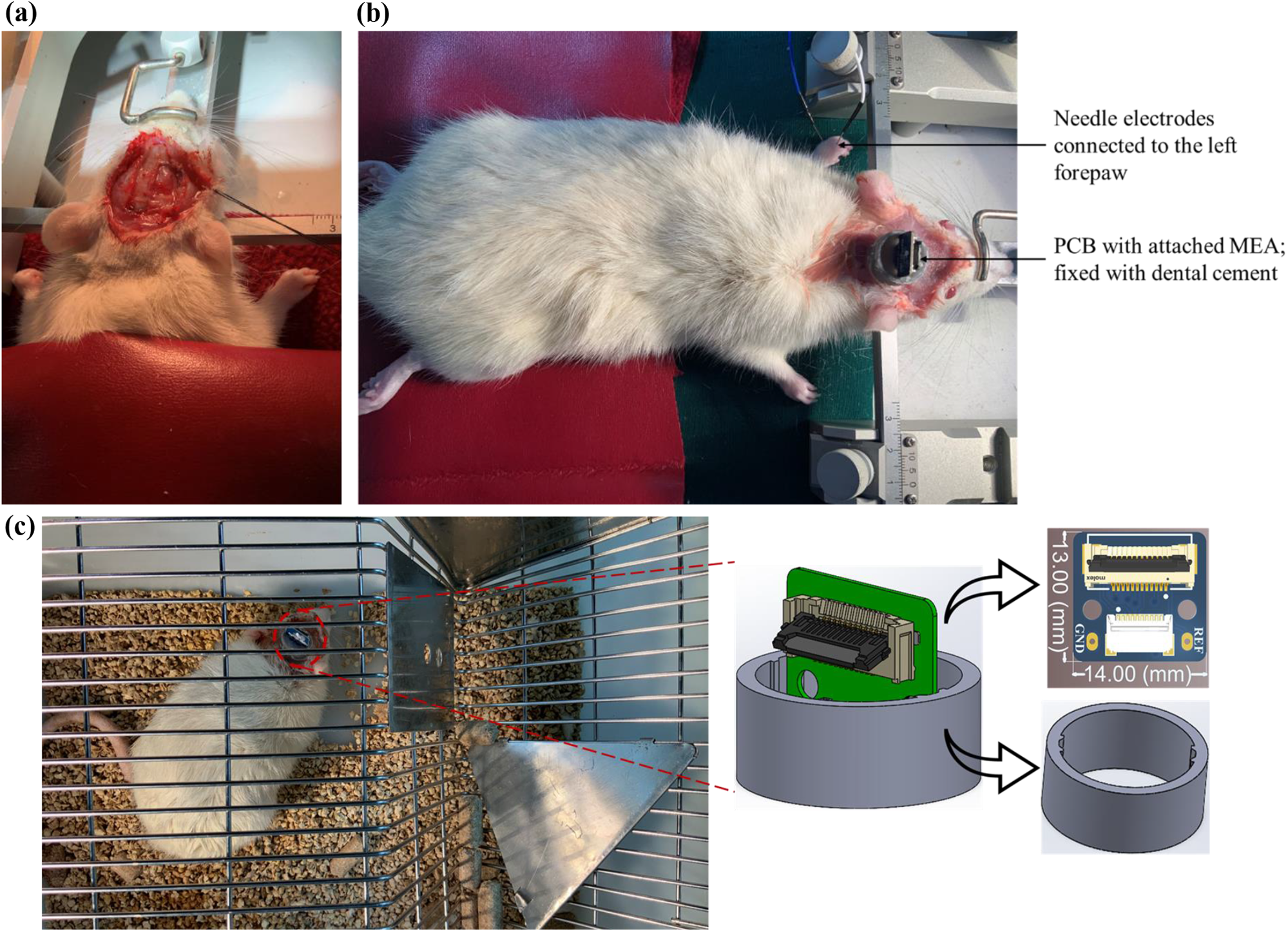
Implantation of the fabricated MEA on the somatosensory cortex. **(a)** Craniotomy of an anesthetized rat to expose the somatosensory cortex of the right hemisphere, **(b)** Sedated rat after implantation of the flexible biodegradable array and the needle electrodes are connected to the left forepaw for stimulation, and **(c)** The rat recovering in a cage after surgery and Design of the 3D printed PCB holder and the PCB.

The rat was administered antibiotics (Enrofloxacin, 10 mg/kg body weight) daily for five days, and a chronic study was performed. The experiments involved recording baseline ECoG activities and applying electrical stimulation to the left forepaw of the rats to induce epileptic activities in the somatosensory cortex, focusing on the somatosensory forelimb (S1FL) area.

The rat was administered Dexmedetomidine and Buprenorphine before applying the electrical stimulation to induce epileptic activities. The Dexmedetomidine was applied at one-hour intervals until the stimulation cycle was over. Initially, the baseline ECoG signals were acquired. Two wire electrodes were connected to the web space of the digits 2/3 and 4/5, and the left forepaw was stimulated by biphasic signals (Figure 3(b)). The peripheral electrical stimulation elicited epileptic activities, and the ictal and interictal episodes were observed. The intraperitoneal administration of an AED, sodium valproate (200 mg/kg body weight), suppressed epileptic activities in the rat. The chronic studies by recording ECoG signals were continued for six days till the signals were recorded faithfully. In chronic studies, the evoked potentials generated by touching the left whiskers resulted in the recording of evoked response by the MEA, ensuring the implantation on the somatosensory cortex. The recorded baselines at an interval of two days and somatosensory evoked responses on the third-day post-surgery were acquired from freely moving rats in a cage (Figure 3(c)). The ECoG signals were recorded by interfacing the acquisition system to the PCB fixed on the head (Figure 3(c) (inset)), with an interval of two days until the recorded signal amplitude was decayed.

The rat was euthanized as per the protocol, and the vital organs, i.e., the brain, liver, kidney, heart, lungs, small intestine, and large intestine, were collected and stored in buffered formalin solution after euthanasia for histological studies.

### 2.8 Electrocorticography signal acquisition

The ECoG signals were acquired from the somatosensory cortex of rats using two OpenBCI Cyton Boards. The biopotentials were acquired by ADS1299 (*Texas Instruments*), an Analog to Digital Converter (ADC) integrated into the board. A dedicated PCB consisting of an intermediate stage of FPC connectors was used to interface the OpenBCI Cyton Boards with the signal channels from the fabricated MEA [22]. OpenBCI Cyton Board, a scientifically proven neural electrical signal acquisition device [23], [24], samples the electrical discharges at 250 Hz, allowing a faithful reconstruction of frequency components up to 125 Hz following Nyquist’s theorem. The boards established synchronous wireless communication to transmit data to computers using Bluetooth Low Energy (BLE) modules.

### 2.9 Signal processing

Custom scripts in MATLAB R2020b (Institutional academic license) with EEGLAB (version 2022.0) were devised for the post-acquisition neural signal analyses [25]. The acquired biopotentials were filtered using a finite impulse response bandpass filter (1 Hz – 100 Hz) [26]. Additionally, the power line interference of 50 Hz was nullified using a notch filter [27]. Similarly, by changing the frequency of a notch filter, power line interference of 60 Hz can be mitigated following electrical standards. Finally, the filtered data was smoothened with a 10-point moving average to remove high-frequency non-neural bursts.

Subsequently, the power spectrum of the preprocessed signals was computed to understand power distribution among different frequency components under three different conditions: (i) Initial baseline, (ii) Epileptiform discharges elicited by peripheral electrical stimulation, and (iii) Recovered baseline after administration of an AED. The power spectrum was calculated by a Welch periodogram following the three-step procedure: (i) Splitting the twenty-second signal into equal, optimal segments with 50% overlapping, (ii) windowing each segment using a Kaiser window, and (iii) averaging the periodograms [28], [29]. Furthermore, the time-frequency analysis was conducted to understand the spatiotemporal changes in filtered signals as a function of time. The spectrogram was approximated by concatenating the short-term Fourier Transform (STFT) of each segment with the Kaiser window, resulting in a matrix. Moreover, the chronic baselines were monitored at regular intervals to observe the effect of MEA dissolution.

## 3. Results and discussion

### 3.1 Studying the degradation of the fabricated MEA

The MEA was fabricated using biodegradable materials, and its degradation was studied in PBS of pH 7.4 at 37 °C with continuous agitation. Figures 4(a) - (d) show the steps of degradation of the encapsulation layer and the metal layer in PBS (volume 30 ml). This dissolution study showed that the encapsulation of the immersed array was degraded in 17 hours, the patterned metal layer was degraded in 21 hours after immersion, and only the substrate remained after 22 hours of immersion. The immersed MEA, MEA after the dissolution of the encapsulation layer, MEA after the dissolution of the patterned metal layer and the encapsulation, and remaining substrate are shown in Figure 4(a), Figure 4(b), Figure 4(c), and Figure 4(d), respectively. This dissolution study shows the degradation of the constituent materials in a bio-mimicked environment at standard physiological temperature.

**Figure 4.**
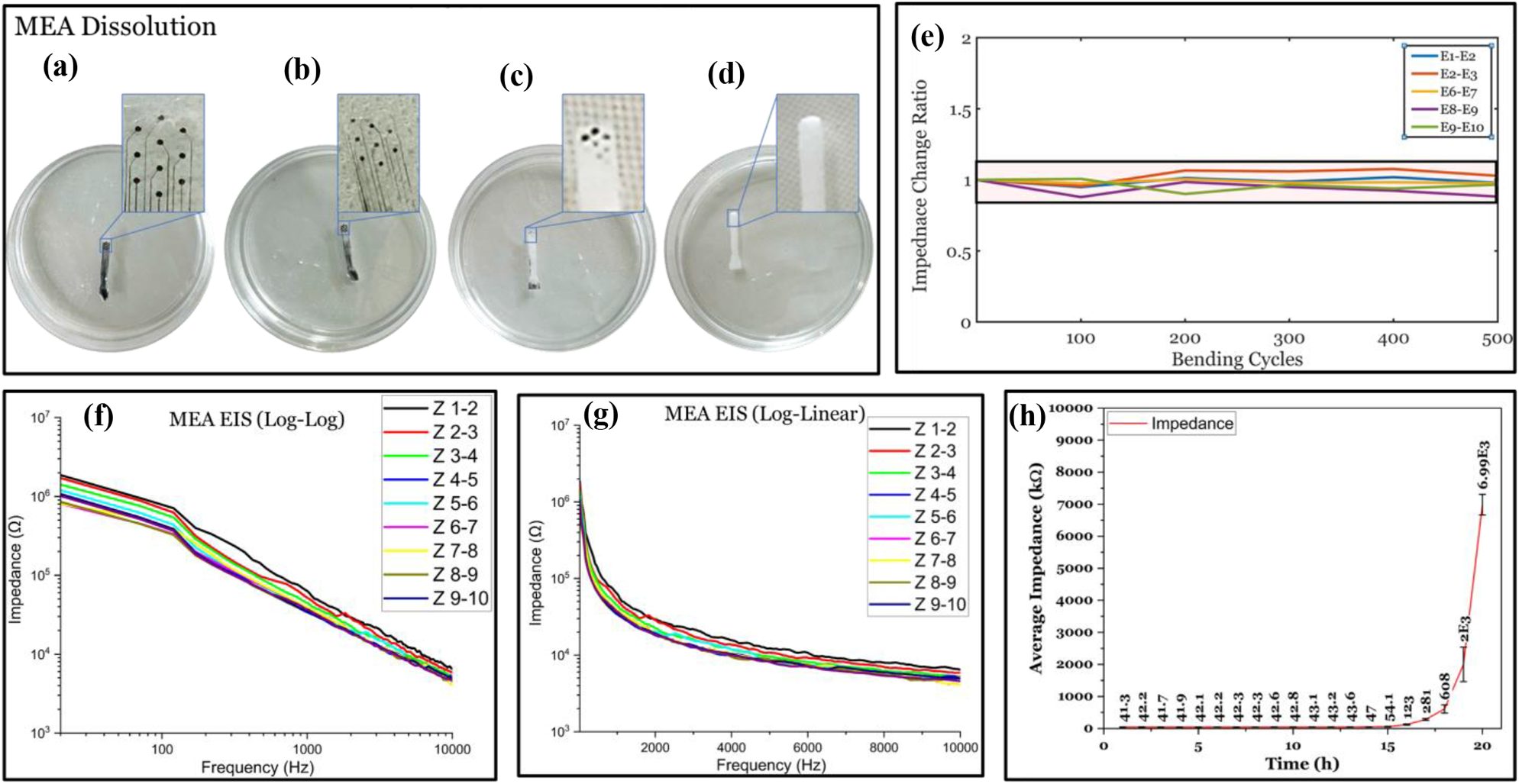
Characterization of the fabricated MEA. **(a)-(d)** Dissolution of the MEA in PBS of pH 7.4: **(a)** MEA after immersion, **(b)** Dissolution of the encapsulation layer of the immersed MEA after 17 hours, **(c)** Dissolution of the electrodes after 21 hours, and **(d)** Dissolution of electrodes and contact pads after 22 hours of immersion. The inset shows the magnified view of the recording electrode array, **(e)** Impedance measurement of five pairs of equidistant electrodes of the MEA for 500 bending cycles to study the change in normalized impedance with bending. The highlighted area shows the change (±10%) in normalized impedance owing to bending; **(f, g)** Electrical Impedance Spectroscopy of the fabricated MEA by measuring the impedance of nine consecutive electrode pairs, and **(h)** The change in average impedance of a pair of consecutive electrodes at 1 kHz during the degradation of encapsulation and the patterned metal layer.

### 3.2 Bending test

The bending test was performed by attaching the MEA to a micromanipulator arm by measuring impedance at 1 kHz after immersing the recording sites in PBS (pH 7.4). The impedance of the electrode pairs was measured from five pairs of equidistant electrodes after 100 bending cycles by 60° using the periodic vertical movement of the micromanipulator up to 500 cycles. The normalized impedance (i.e., 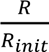, where *R_init_* and *R* denote the initial impedance and the impedance after bending, respectively). These electrode pairs are plotted against the number of bending cycles (Figure 4(e)). Figure 4(e) indicates no significant impedance change because of bending. The impedance of the pairs of equidistant electrodes selected from three columns of the array varies within ±10% (highlighted in Figure 4(e)) of the initial normalized impedance values across the 500 cycles, and the fluctuations are within the expected measurement noise.

### 3.3 Impedance Spectroscopy of the fabricated array

The impedance of nine consecutive electrode pairs was measured by impedance spectroscopy over a frequency range of 20 Hz to 10 kHz with a trigger of 100 mV after immersing the recording area in PBS (pH 7.4) to mimic the *in vivo* condition. The average impedance of the recording electrodes was determined as ∼20 kΩ at 1 kHz, which is in the range for recording ECoG signals [14], [30]. The electrodes had impedances of 20 kΩ ± 3.5 kΩ at 1 kHz. The plotted impedance spectroscopy data is shown in Figure 4(f) and Figure 4(g).

### 3.4 Impedance Spectroscopy to Study Degradation of the MEA

The degradation time of the device was further investigated by studying the impedance spectroscopy of the electrodes during *in vitro* degradation in PBS. The changes in average impedance of the immersed electrode pairs due to degradation were measured at 1 kHz and plotted in Figure 4(h). The initial average impedance of the electrode pairs was ∼40 kΩ with small fluctuations and remained constant for up to 15 hours. Then the average impedance increased drastically after 17 hours and reached ∼2 MΩ after 19 hours of immersion in PBS. The hydrolysis and degradation of the patterned metal layer resulted in this drastic increase in impedance. This study can further confirm the degradation of the encapsulation layer in 17 hours, followed by the degradation of the metal layers; however, the changes in impedance were evident after 16 hours for the immersed MEA.

The gradual increase in electrode impedance (Figure 4(h)) further validated the visual observation of the *in vitro* degradation of the immersed MEA in PBS (Figure 4(a) - (d)). Initially, the standard deviation with the average value of electrode impedance was in the ±4 kΩ range; however, the standard deviation increased drastically after 16 hours due to degradation of the electrodes and the interconnects from recording electrodes to contact pads. Additionally, the drastic change in average impedance shows that this MEA can be used for signal acquisition for 17 hours in this *in vitro* condition.

### 3.5 Acute induction and recording of epileptic activities

The microelectrode array (MEA) was implanted (subdural) on the somatosensory cortex of the right hemisphere to obtain ECoG signals. The screw electrodes drilled on the left hemisphere acted as ground and reference electrodes, with an extended wire to the interfacing unit. After closing the craniotomy defect with dental cement, the implanted MEA was used to record ECoG signals at normal, epileptic, and recovered conditions from three rats (n = 3) to check the reproducibility of the study outcomes. The ECoG traces from all ten channels are plotted for 20 seconds under each neurophysiological condition. The flow of the experiment is shown in Figure 5(a).

**Figure 5.**
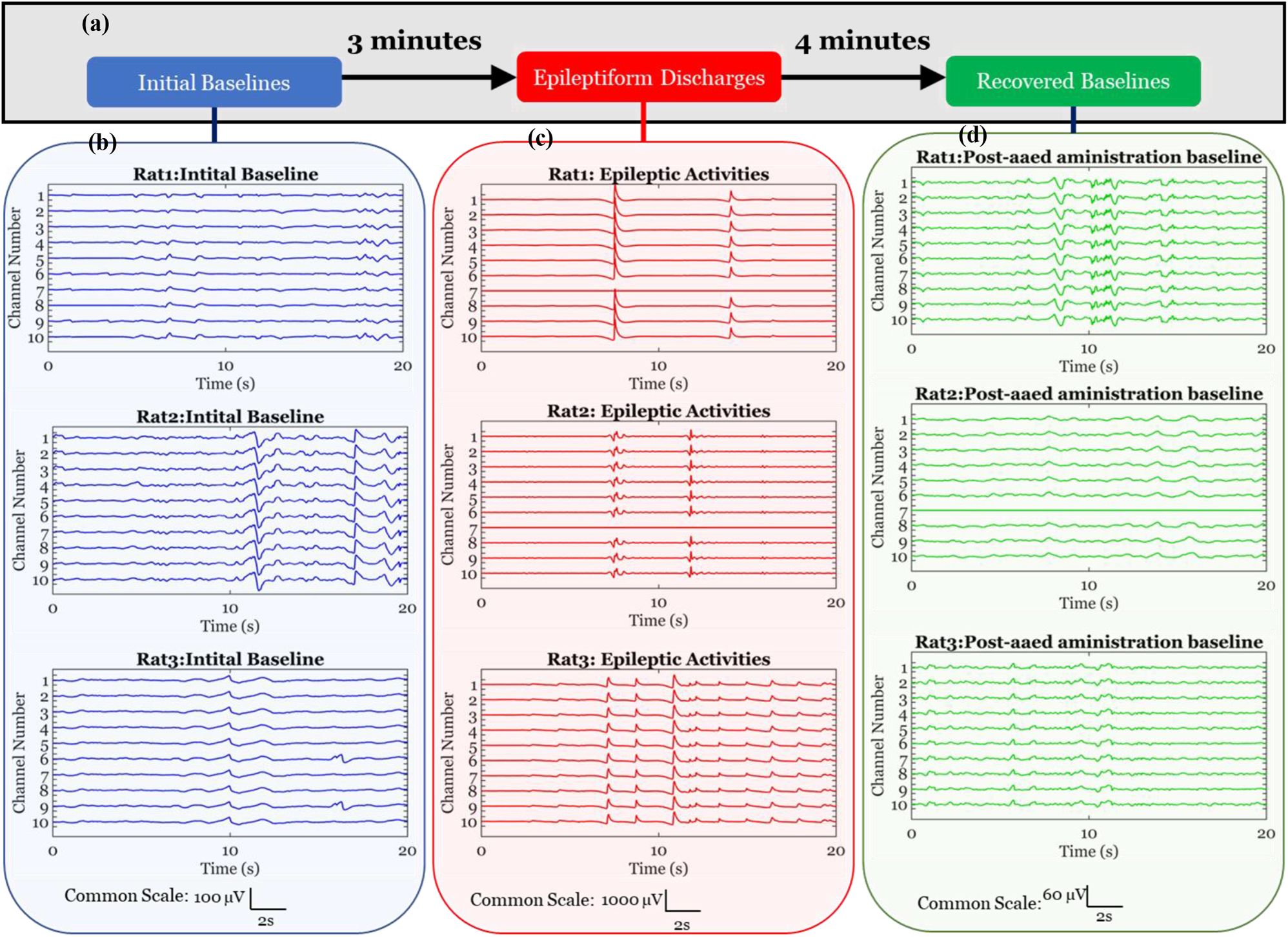
Processed ECoG signals, acquired using fabricated flexible biodegradable MEA. **(a)** Experimental timing diagram, explaining sequential flow with essential timing specifications, **(b)** Baseline ECoG signals (Amplitude Scale: 100 µV), **(c)** Epileptiform discharges induced by peripheral electrical stimulation (Amplitude Scale: 1000 µV), and **(d)** Recovered baselines after AED administration (i.p.) (Amplitude Scale: 60 µV). All these traces were recorded by the MEA and are shown for twenty seconds. Initial baselines, epileptic activities, and recovered baselines are shown at the same scale for all three subjects.

The ECoG signals recorded from the subjects under three neurophysiological conditions are shown in Figure 5(b) - (d). The initial baselines recorded from the cortical surface of the subjects by the MEA are displayed in Figure 5(b), with a blue highlighted region. The baselines exhibited an amplitude in the range of ±50 µV. After acquiring baselines for three minutes, peripheral electrical stimulation (amplitude of 2 mA) was applied to induce epileptiform discharges. The electrical stimulation was applied to the left forepaw periodically for ten runs. The evoked epileptiform activities resulting from the stimulation cycle were exhibited after the 4^th^ run, and similar epileptiform discharges were observed for 3 minutes after the 10^th^ such run. The induced epileptiform discharges are highlighted in the red shaded region in Figure 5(c). These epileptiform discharges have significantly higher amplitude in a range of ±500 µV. These induced activities show repetitive high-frequency discharges, observed by most electrodes. After recording sustained epileptiform discharges, sodium valproate, an antiepileptic drug (AED), was administered intraperitoneally. The recorded ECoG signals exhibited a recovered baseline signal after four minutes of AED administration. Figure 5(d) displays the preprocessed recovered baselines in the green highlight. This shows a low amplitude (±30 µV) baseline, recovered due to sodium valproate (AED) application, confirming that the epileptiform discharges were suppressed. The recorded recovered baselines exhibit a lower amplitude than the initial baselines.

ECoG signals of similar amplitude ranges were obtained from all three subjects, as shown in Figure 5. The electrical stimulation to the forepaw resulted in elicitation of epileptic activities in S1FL area of right hemisphere. As the dimension of the implanted MEA was decided to acquire signals only from this brain region, a change in electrical activities was exhibited by all the electrodes in the MEA.

### 3.6 Time-frequency analysis of recorded ECoG signals

The time-frequency analysis was performed on the recorded ECoG signals under three neurophysiological conditions from the three subjects (n = 3). The signals were recorded from awake rats after administering an analgesic and a sedative. The signals exhibited the presence of similar frequency components and amplitude scales for all subjects. The time-frequency analysis of the signals helps to understand the spectral and temporal features under different neurophysiological conditions.

Temporal traces, power spectrum, and time-frequency analysis of the signals acquired from the first rat are shown for the initial baseline (Figure 6(a1), Figure 6(a2), Figure 6(a3)), induced epileptiform discharges by forepaw stimulation (Figure 6(b1), Figure 6(b2), Figure 6(b3)), and retrieved baseline following intraperitoneal administration of AED (Figure 6(c1), Figure 6(c2), Figure 6(c3)). Striking differences in time-domain amplitude and spectral power were observed between baselines and epileptiform discharges (Figure 6(a1), (b1), and (c1); Figure 6(a2), (b2) and (c2)). The initial baseline (Figure 6(a1)) and recovered baseline (Figure 6(c1)) resemble flat lines as compared to Figure 6(b1), which show epileptic activities in the form of recurrent high-frequency bursts.

**Figure 6.**
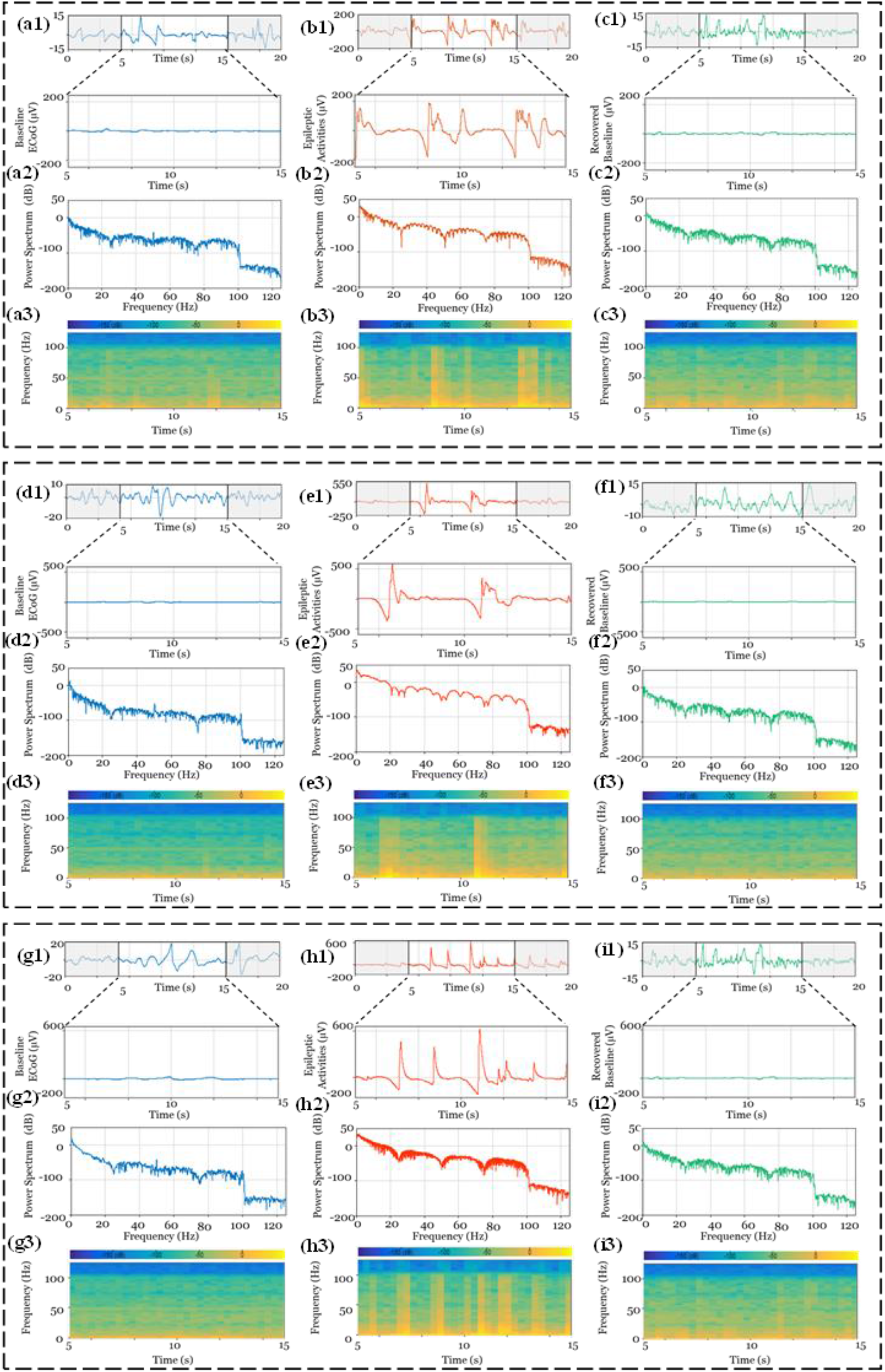
Time-frequency analyses (TFA) of the processed ECoG signals from three rats under three different experimental conditions. **(a,d,g)** TFA of initial baselines, where (**a1,d1,g1)** Recorded baselines, **(a2,d2,g2)** Power spectrums of the recorded baselines, and **(a3,d3,g3)** Spectrograms of the baselines; **(b,e,h)** TFA of induced epileptiform discharges, where **(b1,e1,h1)** Recorded epileptic activities after electrical stimulation, **(b2,e2,h2)** Power spectrums of the epileptic activities, and **(b3,e3,h3)** Spectrograms of the epileptic activities; and **(c,f,i)** TFA of recovered baselines post AED administration, where **(c1,f1,i1)** Recovered baselines after four minutes of AED administration, **(c2,f2,i2)** Power spectrum of the retrieved baselines, and **(c3,f3,i3)** Spectrogram of the recovered baselines.

Moreover, a notable delineation among the baselines and epileptiform discharges was witnessed in time-frequency analysis (Figure 6(a3), (b3), and (c3)), which is more prominent in the frequency range of up to ∼30 Hz. This contrast can be observed by the presence of high amplitude components (represented in yellow colour, Figure 6(b3)) during epileptiform discharges. The dominance of the low-frequency components (up to ∼30 Hz) during induced epileptiform discharges is further confirmed in the power spectrum (Figure 6(b2)) and time-frequency analysis plot (Figure 6(b3)). As a bandpass filter with a range of 1 – 100 Hz was used in preprocessing of the acquired raw biopotentials, the power spectrum and time-frequency analysis results (Figure 6(a2), (a3), (b2), (b3), (c2), and (c3)) showed a sudden decline after 100 Hz.

Time-frequency analysis (Figure 6) shows time domain traces (Figure 6(a1), Figure 6(b1), Figure 6(c1)), power spectrums (Figure 6(a2), Figure 6(b2), Figure 6(c2)), and spectrograms (Figure 6(a3), Figure 6(b3), Figure 6(c3)) at the same scale. Specifically, the identical scale for time domain traces is ± 200 µV, the identical scale for spectrums is +50 dB to −200 dB, and the identical frequency limit for spectrogram is 0 to 125 Hz.

Time-frequency analyses of the processed signals acquired from the second and third subjects are shown in Figures 6(d)-(f) and Figures 6(g)-(i), respectively.

### 3.7 Chronic ECoG Acquisition

The rat was administered antibiotics (Enrofloxacin, 10 mg/kg body weight) daily, and ECoG signals were monitored at an interval of two days. The somatosensory evoked potentials were recorded from the rat on the third-day post-surgery. Touching the left whiskers acted as somatosensory stimulation and was synchronously reflected in the recorded ECoG signals in all electrodes. The obtained traces further ensured the placement of the electrode array on the somatosensory cortical surface. The recorded ECoG signals reflecting the response to touch are shown in Supplementary Figure S1(a)-(c) for all three rats.

The chronic baselines on Day 2, Day 4, and Day 6 showed a visible amplitude decline in all channels, which may be caused by the possible degradation of the electrode array (metal layer) in the *in vivo* condition. Figure 7(a)-(c) shows the ECoG baselines from all three rats on Day 2, Day 4, and Day 6. Several sporadic high-amplitude electrical discharges were observed in some electrodes till the third-day post-surgery; however, these bursts were absent on the sixth day. These observations further confirm the earlier claim of metal layer degradation. The lifetime of such MEAs can be tuned by varying the thickness of encapsulation layer. Thicker the encapsulation layer, more the lifetime of the MEA. Similarly, tuning the ratio of polymers in the composite helps to tune the degradation rate of the encapsulation layer. However, the change in metal layer thickness is not recommended because of change in electrode impedance, which may affect the signal acquisition.

**Figure 7.**
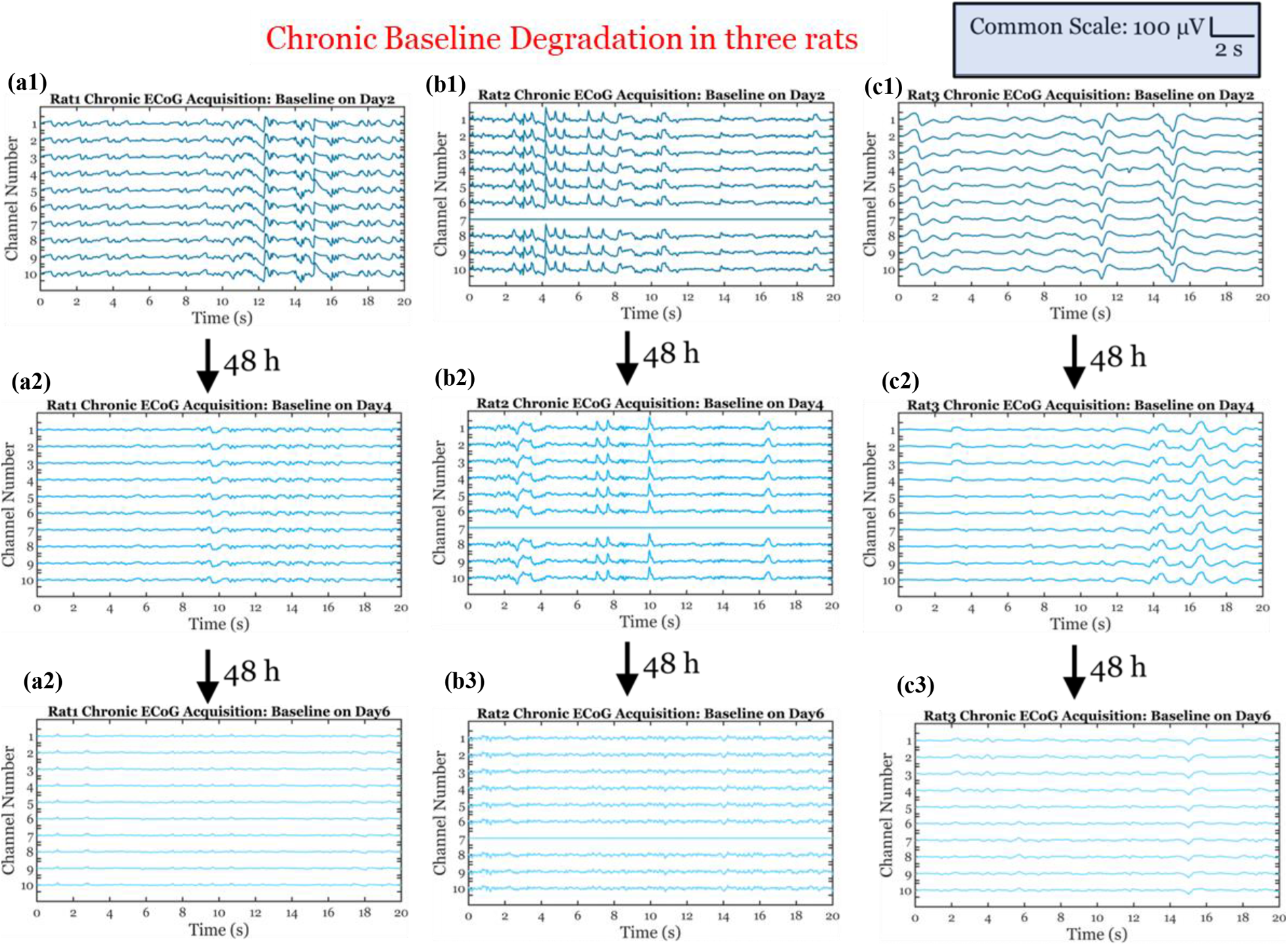
Pre-processed ECoG traces depicting chronic baseline degradation. **(a1-a3)** Rat 1 chronic baselines on Day-2, Day-4, and Day-6 post-surgery, **(b1-b3)** Rat 2 chronic baselines on Day-2, Day-4, and Day-6 post-surgery, and **(c1-c3)** Rat 3 chronic baselines on Day-2, Day-4, and Day-6 post-surgery. The common scale for all nine plots is shown in the top right.

Even though in the *in vitro* dissolution study, the insulation and the metal layers of MEA degraded within 22 hours in PBS (pH 7.4), the *in vivo* experiment shows that the implant (MEA) acquired ECoG for six days. This deviation may be caused by the intracranial CSF volume in rats (∼250 µl) with a continuous flow of ∼2.5 µl/min compared to 30 ml of PBS used for *in vitro* study [31].

Additionally, the degradation of chronic baselines was further confirmed by plotting channel-wise root-mean-square (RMS) value for preprocessed baselines on day 2, day 4, and day 6. Obtained results for three rats are shown in Figure 8(a)-(c).

**Figure 8.**
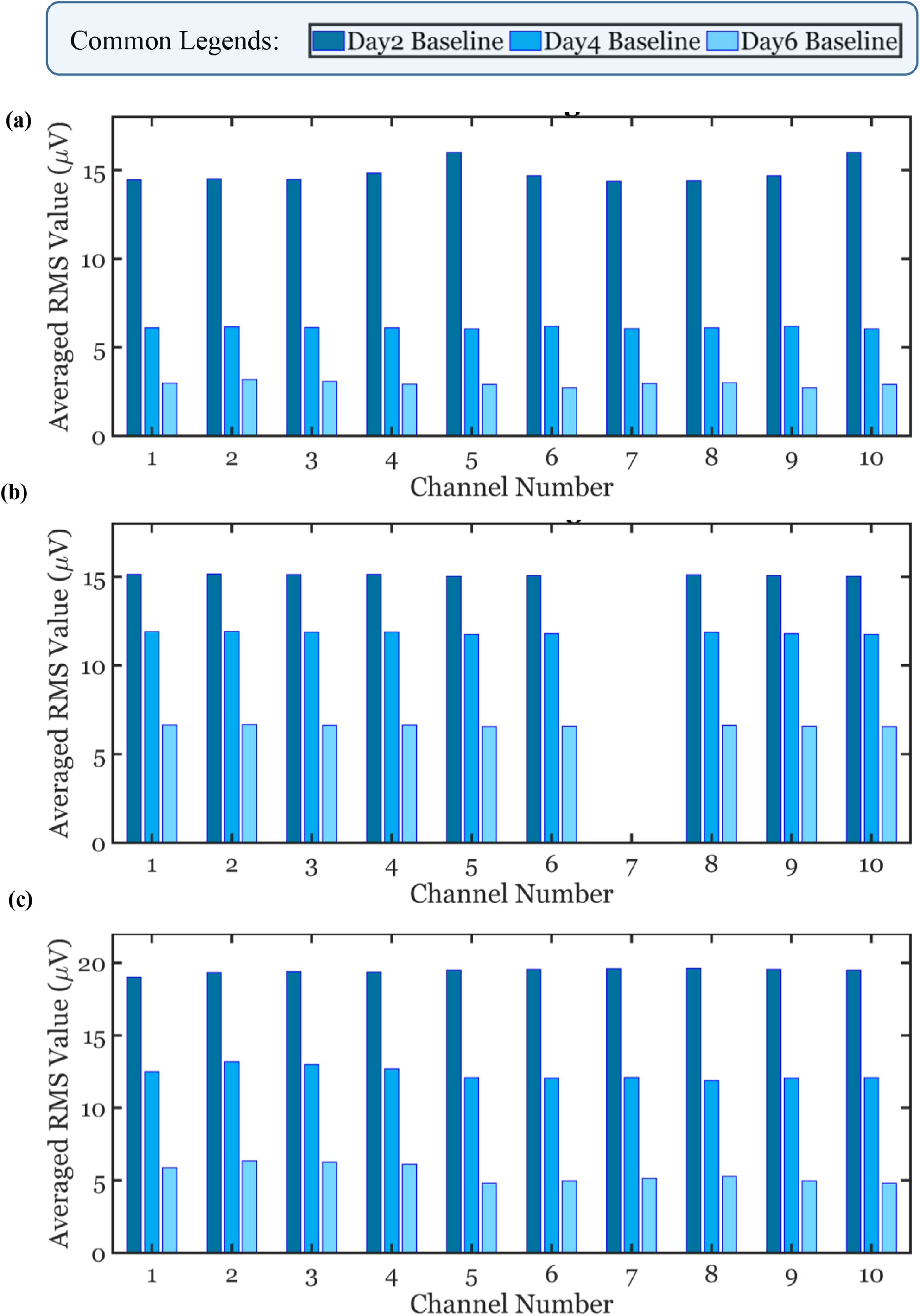
Confirmation of chronic baseline degradation by plotting channel-wise RMS values. **(a)** RMS amplitude degradation in Rat1, **(b)** RMS amplitude degradation in Rat2, and **(c)** RMS amplitude degradation in Rat3. A common colour legend is shown at the top of the figure.

### 3.8 Histological studies of the vital organs

After euthanasia, the vital organs of the rat were recovered and preserved in a 10% neutral buffer formalin solution. The formalin-fixed brain was processed for paraffin embedding, and serial sections of 5 to 7 μm were extracted using a microtome. The brain sections were then mounted on a glass slide, stained with Cresyl Violet, and observed using an optical microscope. Similarly, the kidney, liver, heart, lungs, and intestines were extracted and studied after H&E staining. The histological examinations indicated no tissue damage was observed under a microscope, meaning the implantation did not cause any tissue damage in vital organs. Figure S2 shows the stained sections of the vital organs used for histological studies. Supplementary figure S2(a) shows Cresyl Violet stained extracted rat’s brain sample. Figure S2(b), Figure S2(c), Figure S2(d), Figure S2(e), Figure S2(f), and Figure S2(g) show H&E-stained sections of liver, kidney, heart, lungs, small intestine, and large intestine, respectively.

## 4. Conclusion

The design, fabrication, and characterization of a flexible, biodegradable 10-channel microelectrode array for recording ECoG signals and validation by *in vivo* experiments following MEA implantation have been performed. This novel biodegradable MEA with high electrode density was fabricated using tungsten electrodes and PLLA:PCL (80:20) as the substrate and the encapsulation layer. Before subdural implantation, the fabricated microelectrode arrays were characterized by electrical impedance spectroscopy and *in vitro* degradation. A custom-designed PCB was fixed on the rat’s head and was used to interface the MEA to the acquisition system. An electronic module was developed using a commercially available microcontroller (PSoC^®^ 5LP) for stimulating the left forepaw of the rat to elicit epileptic activities at the somatosensory cortex in the right hemisphere. The sustained epileptiform discharges were suppressed by intraperitoneal administration of sodium valproate, an AED. The somatosensory-evoked responses, epileptic activities, and the recovered baselines were acquired by OpenBCI Cyton Boards. A significant difference in time domain amplitude (seven to ten times) and the presence of low-frequency signals (up to ∼30 Hz) were observed between the epileptiform discharges and the initial and recovered baselines. The *in vivo* degradation of the fabricated array was demonstrated in chronic experiments. These experiments confirmed (i) the competency of the fabricated array to record the ECoG signals at different neurophysiological conditions for a predetermined period and *in vivo* degradation of the array, (ii) elicit of epileptic activities in sedated rats by peripheral electrical stimulation at forepaw without affecting the brain directly by any physical or chemical insult after recovery from ketamine anesthesia, and (iii) competency of the OpenBCI Cyton Board to record brain activities from the cortical surface of a rat to propose this as a low-cost alternative to the commonly used acquisition systems. The histological studies of the vital organs, including the rat’s brain, exhibit an absence of significant inflammation and damage due to chronic implantation. We further envisage that this experimental protocol may help to evaluate and understand the efficacy of AEDs in rat models and the effect of drugs on rat brain activities. It would be interesting to escalate the biodegradable implants as the mainstream implants after validating in larger mammals (e.g., monkeys) and finally, after the human trials with necessary approvals.

## Supplementary Information

Supplementary Information is available with this article.

## Acknowledgements

Hardik J. Pandya acknowledges the partial support from the SERB project (CRG/2020/004427), Abdul Kalam Technology Innovation Fellowship (INAE/121/AKF/49) and DST project (TDP/BDTD/40/2021/General).

## Conflict of interest

The authors declared no conflict of interest concerning with this article.

## Notes

### Competing Interest Statement

The authors have declared no competing interest.

