## Supplementary Information for "Design and fabrication of flexible biodegradable microelectrode array for recording electrocorticography signals"

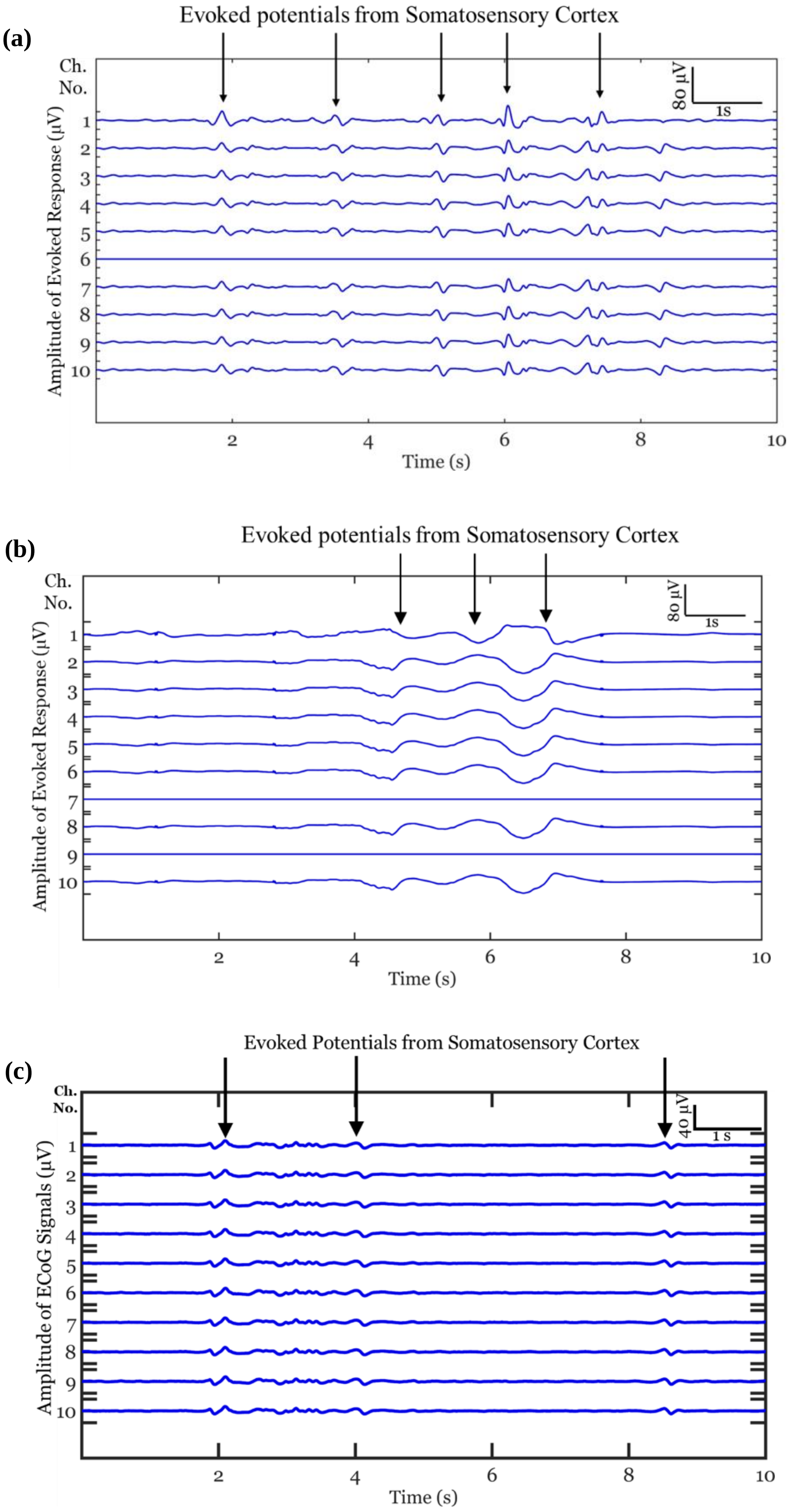


**Figure S1. Recording of somatosensory evoked potentials by the implanted MEA on Day 3 post-surgery, elicited by touching the left whiskers: (a)** Evoked responses in rat 1, **(b)** Evoked responses in rat 2, and **(c)** Evoked responses in rat 3. Scales are shown in right top in each plot. Timestamp for touching whisker is indicated by vertical arrows.

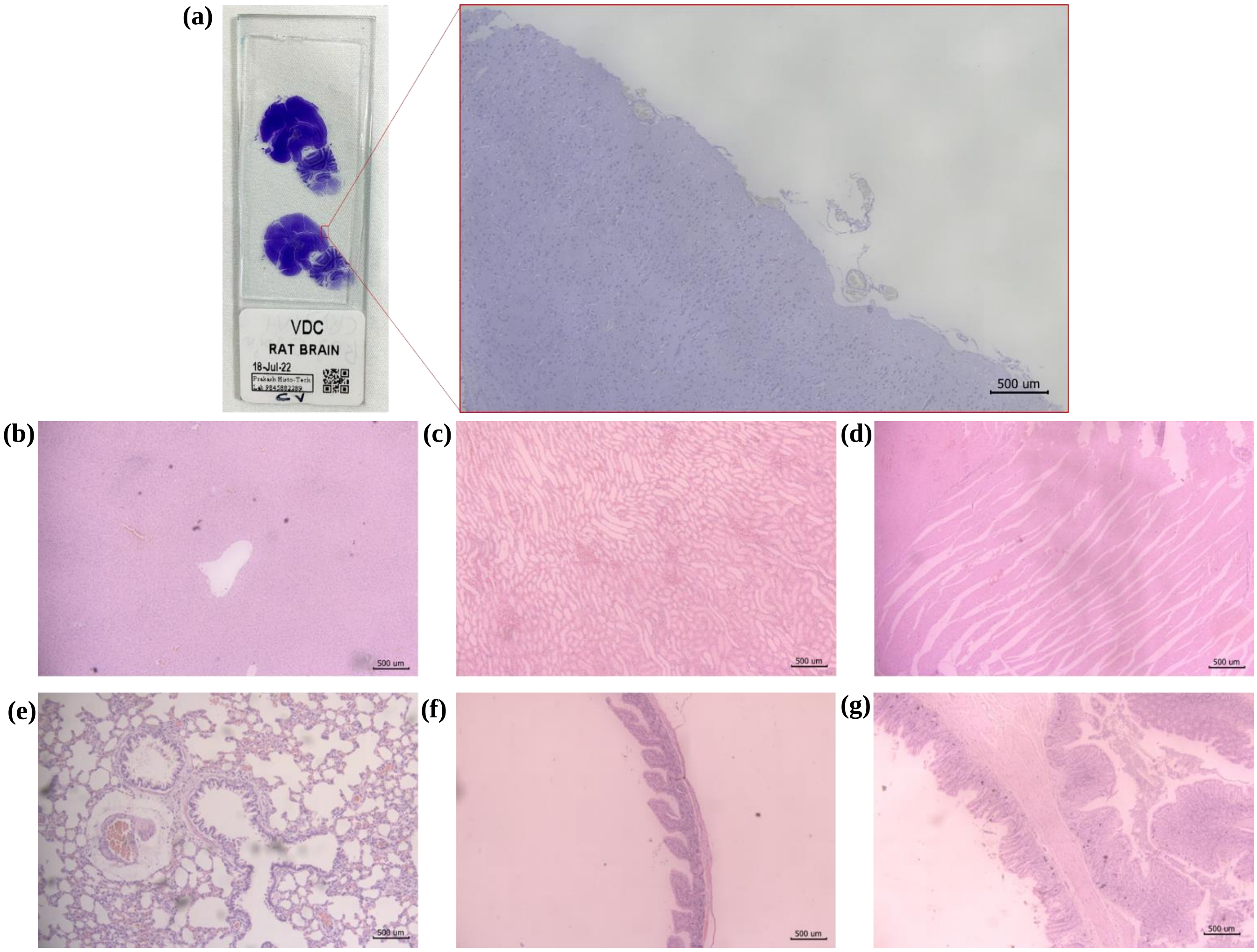


**Figure S2. Histological studies of the vital organs of the rat after chronic experiment: (a)** The slide with Cresyl Violet stained brain section, and (inset) Magnified image of the brain section, captured by an optical microscope for studying the damage due to implantation of the array, **(b)** Magnified image of the section of liver, **(c)** Magnified image of the section of kidney, **(d)** Magnified image of the section of heart, **(e)** Magnified image of the section of lungs, **(f)** Magnified image of the section of small intestine, and **(g)** Magnified image of the section of large intestine.
